# Investigation of a night soil compost psychrotrophic bacterium *Glutamicibacter arilaitensis* LJH19 for its safety, efficient hydrolytic and plant growth-promoting potential

**DOI:** 10.1101/2020.04.24.057588

**Authors:** Shruti Sinai Borker, Aman Thakur, Sanjeet Kumar, Sareeka Kumari, Rakshak Kumar, Sanjay Kumar

## Abstract

Night soil compost (NSC) has traditionally been a source of organic manure in north-western Himalaya. Lately, this traditional method is declining due to modernization, its unhygienic conditions and social apprehensions. Reduction in the age-old traditional practice has led to excessive usage of chemical fertilizers and shortage of water in the eco-sensitive region. Microbiological intervention was attempted to obtain bacterial consortia for accelerated degradation of human faeces in cold climate to improvise this traditional knowledge. *Glutamicibacter arilaitensis* LJH19, a psychrotrophic bacteria was identified as one such potential candidate for the proposed consortia. The bacterium was isolated from NSC of Lahaul valley and exhibited potential hydrolytic activities, the specific activities of amylase, cellulase and xylanase was observed as 186.76 U/mg, 21.85 U/mg and 11.31 U/mg respectively. Additionally, the strain possessed multiple plant growth-promoting (PGP) traits. The bacterium produced 166.11 µg/ml indole acetic acid and 85.72 % siderophore units, and solubilized 44.76 µg/ml phosphate. Whole genome sequence (3,602,821 bps) endorsed the cold adaptation, polysaccharide metabolism, PGP potential of the bacterium. Genome mining revealed biosynthetic gene clusters for type III polyketide synthase (PKS), terpene, and siderophore in agreement with its potential PGP traits. Comparative genomics within the genus revealed 217 unique genes specific to hydrolytic and PGP activity. Negative haemolysis and biofilm production and susceptibility towards all 12 tested antibiotics indicated the bacterium to be a safe bioinoculant. Genomic investigation supported the bacterium safety with absence of any virulence and antibiotic resistance genes. We propose the strain LJH19 to be a potentially safe bioinoculant candidate for efficient degradation of night soil owing to its survivability in cold and its efficient hydrolytic and PGP potential.

## Introduction

The highland agro system of north-western Himalaya lacks productivity and soil fertility due to extreme weather conditions like heavy snowfall, avalanches, landslides, soil erosion, and scanty rainfall (Kuniyal et al., 2004). To increase soil fertility and meet the high demand for manure, traditional winter dry toilets (Fig. 1A) are prevalent in this region. It is a unique traditional method of composting night soil (human excreta) and involves the use of agricultural and household waste locally called ‘*fot*,’ to cover faeces after defecation (Fig. 1B) (Oinam *et al*., 2008). The resultant manure is supplemented in the fields to sustain the agro-ecosystem (Indian science wire, 2019; Oinam *et al*., 2008). However, due to the effect of modernization and the introduction of modern septic toilets, the practice of dry toilets and night soil composting is nearing demise. Other factors such as foul odour, availability of subsidized chemical fertilizers, rise in the standard of living, the difficulty of finding labour and social apprehensions have also contributed to the decline of the age-old practice (Oinam et al., 2008). For decades this age-old practice of night soil compost (NSC) have conserved water during severe winter and the organic manure formed have sustained the agro-ecosystem. Now due to abandoning this practice, there is excessive use of chemical fertilizers and shortage of water in the region for agriculture purpose. Maintaining the agro-ecosystem, conserving water and promoting dry toilets in the region is of utmost priority. It was suggested to combine the scientific knowledge with traditional dry toilets so that the practice becomes safer and hygienic (Oinam et al., 2008). However, we failed to retrieve any literature support on any scientific intervention addressing this problem.

**Fig. 1.**
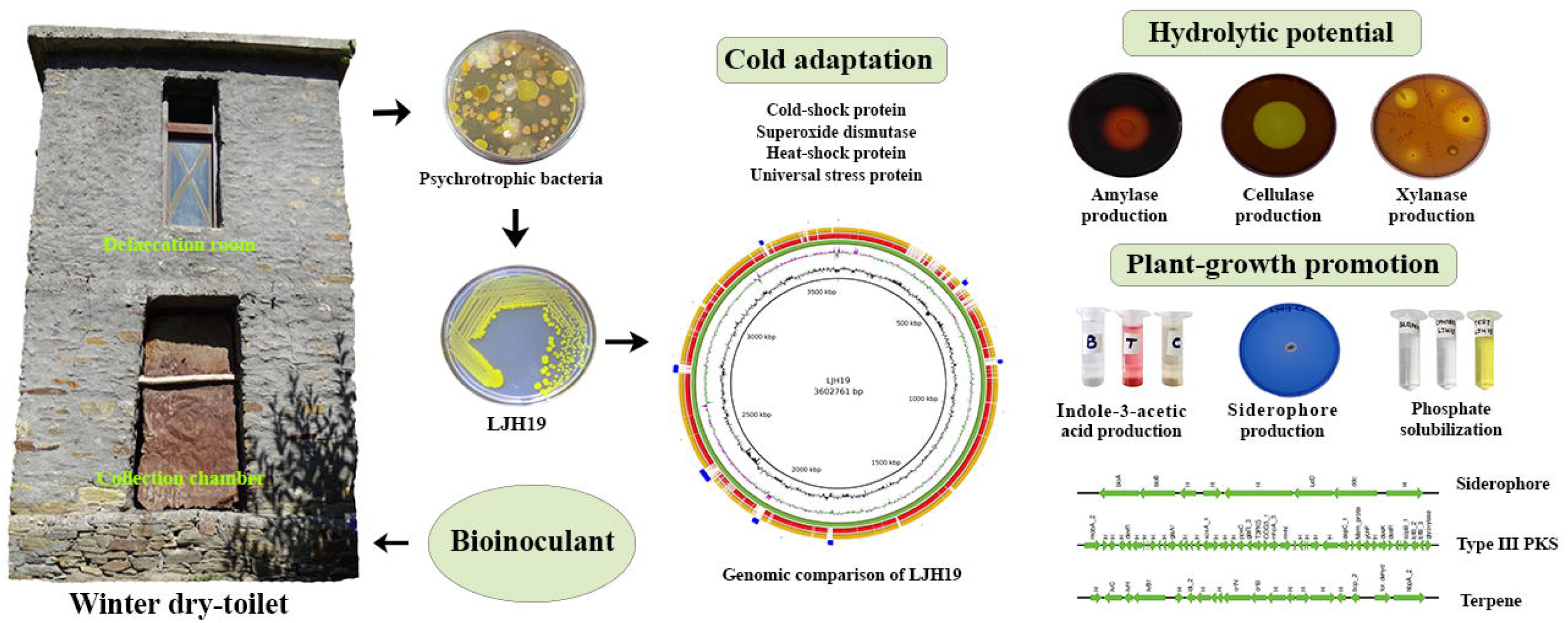
Traditional winter dry toilet of Lahaul valley: **A)** Traditional winter dry toilet attached to the living room of the main house. The upper storey is used for defaecation, while lower storey is a collection chamber where composting takes place. **B)** Inside view of defaecation room. After defaecation people cover the faces with *fot* (dry cattle/sheep dung, kitchen ash, dry grass/leaves). C) Collection of night-soil compost (NSC) sample from the collection chamber. D) NSC pile dumped in open fields for further curing.

Microbial involvement for rapid decomposition in a cold climate (below 20°C) using psychrotrophic bacterial consortium has been emphasized earlier (Hou et al., 2017). Here, a psychrotrophic bacterial community weighs special attention to accelerate the biodegradation process in night soil composting. The lower microbial load in the initial composting process delays the mesophilic phase due to slower biomass degradation resulting in a shorter thermophilic phase (Hou et al., 2017). These conditions subsequently affect the maturity of compost and risks safety and contamination (Millner et al., 2014).

Plant growth-promoting bacteria (PGPB) also plays an important role in maintaining soil fertility. It directly stimulates plant growth by increasing the availability of the nutrients, often found in the form which is unutilized by the plants such as iron, nitrogen, and phosphorous (de Souza et al., 2015). PGPB also produces phytohormones (indole acetic acid), growth regulators (siderophores) and solubilise phosphate to modulate plant growth and development (Numan et al., 2018). The use of PGPB thus gains importance, for sustenance and supporting the agro-ecosystem of Lahaul valley.

Pathogenicity was the main concern while exploring night-soil compost (NSC) for potential psychrotrophic bacteria. Human faeces are known to contain large amounts of enteric microorganisms and many opportunistic pathogens (Heinonen-Tanski and van Wijk-Sijbesma, 2005). The low ambient temperature of the Lahaul region and lower metabolic activity of microbial load might increase the possibilities of poor thermal inactivation of pathogenic strains in NSC. Such a scenario promotes the chances of low-level contamination of pathogenic strains. Apparently, this expresses the worry about the potential of isolated bacterial strains to have harmful impacts on the human, environment, and crop health (Avery et al., 2012).

In the course of finding potential non-pathogenic, hydrolytic and plant growth-promoting (PGP) psychrotrophic bacteria from NSC, we obtained a bacterial strain LJH19 that showed remarkable PGP traits and considerable capabilities of hydrolytic activity in *in-vitro* assays. Owing to its cold adaptation, efficient hydrolytic activity, PGP potential, and origin from faecal compost, whole-genome sequencing was performed to elucidate the genetic basis of the catabolic activities, PGP traits, and analysis for pathogenicity determinants. Further, to explore the habitat-specific gene repertoire, we performed comparative genomics of LJH19 with all the available strains of same genus. On the basis of a unique genome region across the strain LJH19, a comparison was withdrawn with the closely related strains. Biosynthetic gene clusters in the genome of LJH19 were also identified and further to evaluate bacterial safety, the presence of antibiotic resistance gene cluster across all the strains was assessed. To the best of our knowledge, this is first such a report from NSC of north-western Himalaya on bacterial safety and its functional characteristics.

## Materials and method

### Sampling source, strain isolation and hydrolytic potential

NSC samples were collected from the collection chamber of night-soil composting toilet, locally termed as “Ghop” of Jundah village (32.64°N 76.84°E) of Lahaul valley (Fig. 1C). The samples were collected from the core of the compost pile in sterile plastic bottles using stainless steel spatula in triplicates and stored in ice box containing ice packs. The samples were then immediately transported to the laboratory, and processed. The temperature was noted at the time of sampling itself by inserting the handheld digital thermometer (model: ST9269, MEXTECH, India) into the core of the compost pile. Air dried solid sample was mixed with milli-Q at a ratio of 1:10 vortexed and were kept overnight to check pH and electrical conductivity (EC) using digital pH and EC meter (Eutech, India). For the analysis of total available nitrogen, phosphorus and potassium, the samples were dried at room temperature, finely grounded and sieved. All the chemical analysis was performed as per the standard methods for testing of compost materials (TMECC, 2002).

The bacterial strain LJH19 was isolated from NSC samples while screening for potential psychrotrophic hydrolytic bacteria. The isolation was carried out using serial dilution method and spread plate method on nutrient agar (NA) medium (HiMedia) at 15°C. The bacterial isolation was performed in the Class II, Type A2 Biological Safety Cabinet (Thermo Scientific™). The optimum growth conditions of the LJH19 strain were determined by incubating the culture at various temperatures (4-50°C), NaCl concentration (1-10%), and pH (2 to 10) range. The production of hydrolytic enzymes by the LJH19 strain was initially screened using a plate assay method. The exponentially grown culture of LJH19 was spot inoculated on Carboxymethylcellulose (CMC) agar (Kasana et al., 2008), Starch agar (Hi-Media), Xylan agar (Alves-Prado et al., 2010), Tributyrin agar (Hi-Media) plates and incubated at 15°C for 48 hrs. The clear halo zones around the colony indicated positive results. The enzymatic index (EI=Diameter of the halo of hydrolysis/Diameter of the colony) was calculated as described previously by Vermelho and Couri, 2013. For quantitative estimation of polysaccharide degrading enzymes viz. cellulase, xylanase, and amylase, the microplate-based 3, 5-dinitrosalicylic acid colorimetry method was followed using 1% (w/v) carboxymethylcellulose (CMC), 1% birchwood xylan (HiMedia) and 1% soluble starch (HiMedia) as the substrate (Xiao et al., 2005).

### Haemolysin and protease assay, biofilm formation and antibiotic susceptibility profile

For assessment of pathogenic potential, we assayed LJH19 for protease and haemolysin activity using a plate assay method (Igbinosa et al., 2017). The strains *Staphylococcus aureus* subsp. aureus (MTCC 96), *Bacillus subtilis* (MTCC121), *Escherichia coli* (MTCC 43), *Micrococcus luteus* (MTCC 2470) were used as a positive control for haemolytic activity. Haemolytic activity was interpreted according to Buxton, (2016).

Biofilm formation was evaluated according to Basson and Igbinosa (Basson et al., 2008; Igbinosa et al., 2017).) with slight modifications. The cells adhered were stained with 200 μL of 0.5% crystal violet for 10 min. The optical density (OD) readings from respective wells were obtained at 595 nm. The cutoff OD (ODc) for the test was set using the formula (Mean OD of negative control + 3x Standard deviation) and results were interpreted as previously described by Basson et al., (2008). The wells containing only TSB broth (200 μL) served as negative control while the well-containing strains Staphylococcus aureus subsp. aureus (MTCC 96), Bacillus subtilis (MTCC121), Escherichia coli (MTCC 43), Micrococcus luteus (MTCC 2470) was used as a positive control. The test organisms were characterized as non-biofilm producers (OD<ODC), weak (ODC < OD< 2ODc), intermediate (2ODc < OD< 4ODc), and strong (OD>4ODc).

Antibiotic susceptibility profiling was carried out by using the Kirby-Bauer method (Bauer et al., 1966). The antibiotic discs (HiMedia) used were 15 mcg, Azithromycin (AZM); 10 mcg, Ampicillin (AMP); 5 mcg, Ciprofloxacin (CIP); 30 mcg, Chloramphenicol (CHL); 15 mcg Erythromycin (E); 10 mcg, Gentamycin (G); 30 mcg, Kanamycin (K); 10 Units, Penicillin-G (P); 5 mcg, Rifampicin (RIF); 10 mcg, Streptomycin (S); 30 mcg, Tetracycline (TE); 30 mcg, Vancomycin (VA). The plates were incubated at 15°C for 48 hrs. Zones of clearance was measured in millimetre (mm) and interpreted as Resistant, Intermediate or Sensitive using guidelines provided by the manufacturer.

### Plant growth-promoting (PGP) attributes

Indole acetic acid (IAA) production by LJH19 was studied according to Goswami et al., (2014) by supplementing the nutrient broth (100 ml) with L-Trytophan (200 μg ml−1). Siderophore production by LJH19 was initially screened on Chrome Azurol Sulphonate (CAS) agar at 15°C (Lynne et al., 2011). Quantitative estimation of siderophore was done using CAS-shuttle assay (Goswami et al., 2014) by growing LJH19 in iron-free CAS-broth (pH 6.8) at 15°C at 150 rpm. Ammonia production was quantified spectrophotometrically (Cappuccino. and Sherman, 2014). LJH19 was grown in peptone water at 15°C for 10 days at 150 rpm. Inorganic phosphate solubilisation was estimated by the vanado-molybdate method (Gulati et al., 2010) using NBRIP broth containing 0.5% tricalcium phosphate (TCP).

### Strain identification, phylo-taxono-genomics, and gene content analysis

The genomic DNA was extracted using the conventional CTAB method (William et al., 2004) and for identification, partial 16S rRNA gene sequencing of V1 and V9 regions using 27F and 1492R primers, was performed. To provide a genetic basis to the experimental evidence, we performed whole-genome sequencing using PacBio RS-II (Pacific Biosciences, US) as previously described (Himanshu et al., 2016; Kumar et al., 2015). The draft genome sequence was deposited in NCBI GenBank with accession number SPDS00000000. Strain identification was done at the species level using EzTaxon (https://www.ezbiocloud.net/identify). Manual curation of the genomes from a public repository for the closest match was performed with NCBI Genome (https://www.ncbi.nlm.nih.gov/genome/?term=Glutamicibacter). Genome quality was assessed using CheckM v1.1.2 (Parks et al., 2015) in terms of its completeness and contamination present.

Strain phylogeny was assessed using the 16S rRNA gene sequence as well as whole-genome phylogeny using PhyloPhlAn v0.99 (Segata et al., 2013). For the construction of the 16S rRNA gene phylogenetic tree, all the full-length 16S rRNA reported strains of genus *Glutamicibacter* were used. *Micrococcus luteus* (Nucleotide accession No: AF542073) was used as an outgroup. The 16S rRNA gene phylogenetic tree was constructed using ClustalX v2.1 (Larkin et al., 2007) and FastTree v2.1.8 (Price et al., 2010) with default parameters. Whole-genome phylogeny was constructed using PhyloPhlAN (It uses a 400 most conserved gene across bacterial domains and constructs its phylogeny). All the genomes (Table 2) were annotated using prokka v1.14.6 (Seemann, 2014). The orthoANI v1.2 (Yoon et al., 2017) was performed to infer the taxonomic relatedness of the strain LJH19. ANI value matrix obtained was used for generating heatmap using the webserver of Morpheus (https://software.broadinstitute.org/morpheus). Further, 10 strains forming a clade were considered for pan-genome analysis with a 95% cutoff using Roary v3.6.0 (Page et al., 2015). The unique gene present in the strain LJH19 was fetched and annotated with eggNOG mapper v1 (Powell et al., 2014) (http://eggnogdb.embl.de/app/home#/app/home). Chromosomal maps (Alikhan et al., 2011) for comparison of the closely related strains and visualization of the unique genomic region across the strain LJH19 to the type strain RE117 and JB182 are marked in the figure (Fig. 3).

To identify the biosynthetic gene clusters (BGCs) in genome *G. arilaitensis* LJH19, we used a web-based server of antiSMASH v5.0 (https://antismash.secondarymetabolites.org/#!/start) (Blin et al., 2019) and cluster image of the identified biosynthetic gene was prepared with EASYfig v2.2.2 (Sullivan et al., 2011). To assess the presence of the antibiotic gene cluster across the strains, a web-based server of Resistance gene identifier (RGI) v5.1.0 module of Comprehensive Antibiotic Resistance Database (CARD) v3.0.8 (Alcock et al., 2020) was used with strict mode. The pathogenic potential of LJH19 was also assessed using the PathogenFinder web service under automated mode (Cosentino et al., 2013).

## Results and Discussion

### Physico-chemical properties of NSC samples and bacterial characterisation

Sampling in triplicate was conducted from the collection chamber of the traditional night soil toilet “*ghop*”. The temperature of the sample was noted to be 9.9°C at the core of the pile. The pH value of the compost sample was observed to be 10, while the electrical conductivity (EC) noted was 1674 µS. The available nitrogen, phosphorous and potassium in the NSC sample was 2297.6 ± 99.4 ppm, 117.11 ± 0.34 ppm, and 22534.11 ± 73.08 ppm respectively.

In an attempt to explore the bacterial diversity of NSC, we isolated 130 bacterial strains belonging to different phyla (unpublished data). In this study, the hydrolytic potential of an opaque, yellow-pigmented bacterium LJH19 was determined on agar plates (Table 1) which showed significant activity for different substrates viz. corn starch, CMC and birchwood xylan (Fig. S1) (Table 1). To further characterize this bacterium, partial 16S rRNA gene sequencing was performed, which gave best hit with *G. arilaitensis* Re117 with 100% identity and coverage of 96.5% in EzTaxon Biocloud (https://www.ezbiocloud.net/identify). The 16S rRNA partial gene sequence was deposited in the NCBI through BankIt web-based sequence submission tool under accession number MT349443.

**Table 1.**
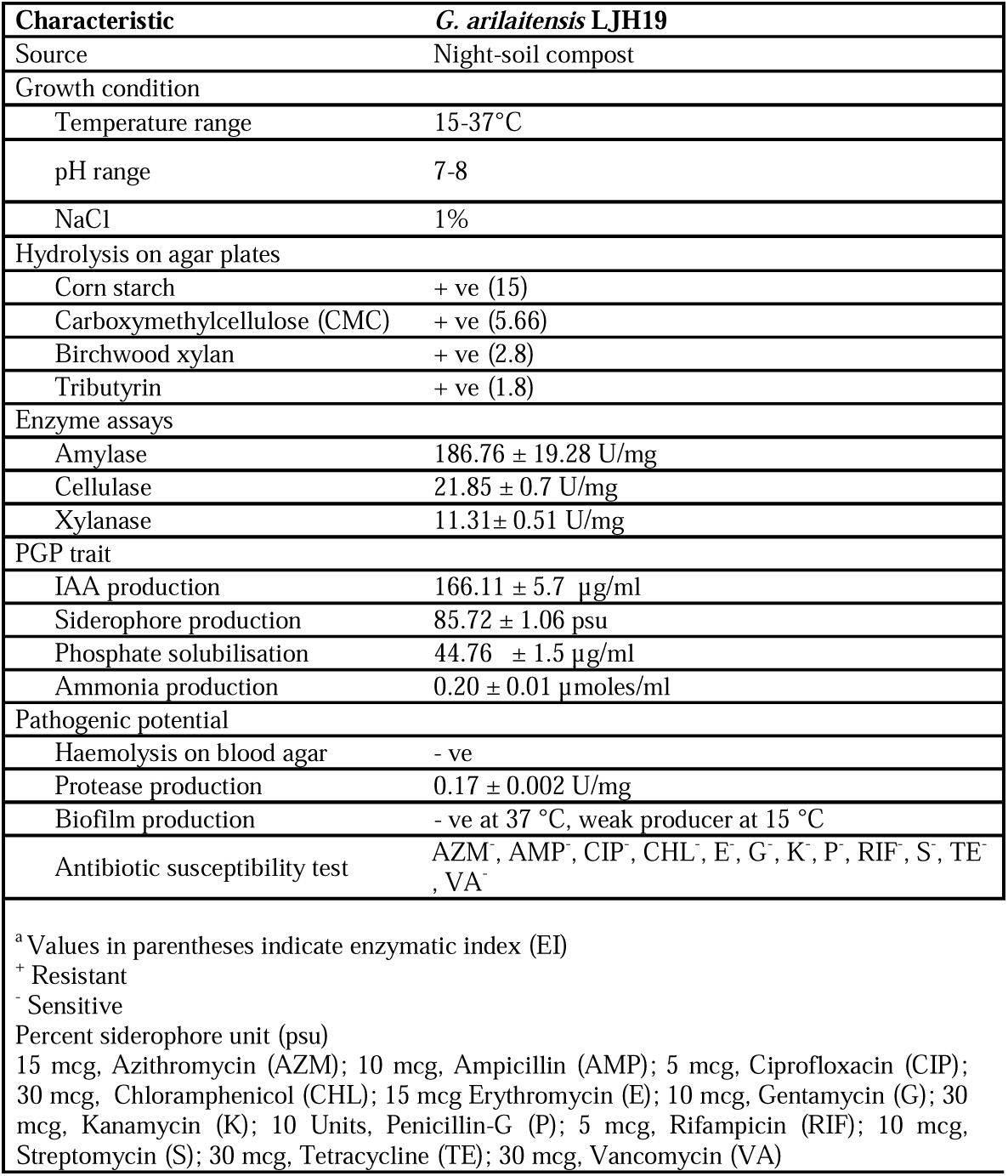
Physiological characterization, hydrolytic, plant growth promoting, and pathogenic attributes of *G. arilaitensis* LJH19

**Table 2:** Genome features of all the strain of *Glutamicibacter* sp. and strain LJH19 with its geographical attributes.

| Genome Name | Genome Status | Coverage (x) | # Contigs | Genome Length (bps) | % GC Content | # RefSeq CDS | Isolation source | %Completeness/<br>%Contamination | Accession No. |
| --- | --- | --- | --- | --- | --- | --- | --- | --- | --- |
| *Glutamicibacter arilaitensis* LJH19 | WGS | 153.0x | 4 | 3602821 | 59.60 | 3245 | Night soil compost | 99.65/1.38 | NZ_SPDS00000000 |
| *Glutamicibacter arilaitensis* Re117 | Complete | NA | 2 | 3909664 | 59.28 | 3423 | Cheese | 99.74/0.46 | NC_014550 |
| *Glutamicibacter soli* M275 | WGS | 500.0x | 319 | 3965931 | 64.09 | 3800 | - | 99.28/2.14 | NZ_WYDN00000000 |
| *Glutamicibacter soli* NHPC-3 | WGS | 100.0x | 25 | 3840670 | 64.34 | 3495 | Tea plant rhizospheric soil | 99.28/0.69 | NZ_POAF00000000 |
| *Glutamicibacter uratoxydans* NBRC 15515 | WGS | 152.0x | 49 | 3786014 | 61.09 | 3533 | - | 99.74/0.58 | NZ_BJNY00000000 |
| *Glutamicibacter nicotianae* NBRC 14234 | WGS | 162.0x | 43 | 3554887 | 61.92 | 3311 | - | 99.74/1.19 | NZ_BJNE00000000 |
| *Glutamicibacter nicotianae* OTC-16 | Complete | 380.0x | 3 | 3797724 | 61.72 | - | Active sludge around pharmaceutical factory | 99.28/1.53 | NZ_CP033081 |
| *Glutamicibacter* sp. JC586 | WGS | 100.0x | 28 | 3524842 | 55.57 | - | Soil | 99.51/1.49 | NZ_VHIN00000000 |
| *Glutamicibacter mysorens* DSM 12798 | WGS | 34.0x | 1 | 3459735 | 61.95 | 3177 | - | 99.74/0.96 | NZ_PGEY00000000 |
| *Glutamicibacter arilaitensis* JB182 | WGS | 50.0x | 12 | 3947886 | 59.16 | 3708 | Cheese rind | 99.74/0.54 | NZ_PNQX00000000 |
| *Glutamicibacter* sp.<br>HZAU | WGS | 100.0x | 40 | 3508313 | 61.70 | 3212 | Bto:0003809 | 99.68/0.38 | NZ_SCKZ00000000 |
| *Glutamicibacter* sp.<br>V16R2B1 | WGS | 12.0x | 86 | 3479500 | 65.68 | 3197 | Date palm rhizosphere | 99.05/1.15 | NZ_VATX00000000 |
| *Glutamicibacter* sp.<br>BW80 | WGS | 50.0x | 78 | 4049569 | 60.38 | 3720 | Cheese rind | 99.74/0.61 | NZ_NRGV00000000 |
| *Glutamicibacter* sp.<br>BW78 | WGS | 50.0x | 43 | 3419483 | 64.04 | 3133 | Cheese rind | 99.77/0.61 | NZ_NRGU00000000 |
| *Glutamicibacter* sp.<br>BW77 | WGS | 50.0x | 90 | 3902028 | 56.57 | 3605 | Cheese rind | 99.51/0.5 | NZ_NRGT00000000 |
| *Glutamicibacter*<br>*halophytocola*<br>DR408 | Complete | 231.0x | 1 | 3770186 | 60.19 | 3384 | Rhizosphere of<br>Soybean | 99.51/1.15 | NZ_CP042260 |
| *Glutamicibacter* sp.<br>0426 | WGS | 241.0x | 23 | 3549469 | 62 |  | Soil | 99.74/0.8 | NZ_MPBI00000000 |
| *Glutamicibacter*<br>*creatinolyticus*<br>LGCM 259 | Complete | 286.0x | 1 | 3309128 | 65.55 | 2882 | Abcess of a mare | 99.51/0.69 | NZ_CP034412 |
| *Glutamicibacter*<br>*mysorens* NBRC<br>103060 | WGS | 130.0x | 15 | 3427456 | 62.02 | - | - | 99.74/0.96 | NZ_BCQO00000000 |
| *Glutamicibacter* sp.<br>ZJUTW | Complete | 100.0x | 2 | 3673306 | 61.79 | 3379 | Activated sludge | 99.74/0.8 | NZ_CP043624 |

### Hydrolytic activity and plant growth promoting attributes of LJH19

The biodegradation of complex polysaccharide by bacteria requires a cocktail of enzymes to depolymerize it to oligosaccharides and monomer sugars (Awasthi et al., 2015). Amylases plays a crucial role in hydrolysis of starchy molecules which specifically acts on alpha-1,4-glycosidic linkages to yield maltose, D-glucose, dextrins and shorter oligosaccharides (Jurado et al., 2015). Quantitatively, LJH19 showed the production of amylase enzyme with specific activity of 186.76 ± 19.28 U/mg (Fig. S2) using corn starch as substrate. Similarly, LJH19 also exhibited the production of cellulase enzyme with specific activity of 21.85 ± 0.7 U/mg (Fig. S2) using CMC as a substrate at 15°C. Cellulase enzyme is involved in hydrolysis of cellulose and hemicellulose, which are the major constituent of plant cell wall (Kostylev and Wilson, 2012). Previous studies (Aarti et al., 2018; Aarti et al., 2017) also found that *G. arilaitensis* strain ALA4 exhibit efficient amylase and cellulase activity. This evidence supports the capability of LJH19 to produce the amylase and cellulase enzyme.

Biodegradation of hemicelluloses is attained by the collective action of a variety of hydrolytic enzymes (Chandra et al., 2015). Since xylan is the major polysaccharide found in hemicelluloses, xylan β-(1, 4)-xylosidase plays a key role in its degradation. In addition to cellulase, LJH19 also showed the production of xylanase enzyme with specific activity of 11.31± 0.51 U/mg (Fig. S2) using birchwood xylan as a substrate at 15°C. Based on these results LJH19 presents the considerable potential of hydrolysing complex polysaccharides (starch, cellulose and xylan) which are the major constituents of residues used in NSC while surviving under low ambient temperature. It is critical for a bacterium to thrive in cold regions as well as to perform collectively, ensuring efficient composting (Hou et al., 2017).

During the process of decomposition, a large number of nutrients are released (Biswas and Kole, 2017). These nutrients are lost from agricultural systems due to leaching, surface runoff and eutrophication, eventually remaining unavailable for plant uptake (Adesemoye and Kloepper, 2009). The plant growth-promoting bacteria (PGPB) aids in improving nutrient uptake and increases the efficiency of applied compost. It plays a pivotal role as biofertilizers since they promote plant growth in a nutrient-limited soil by improving the nutrient availability, and phytohormone production (Dey et al., 2004). Since, soil temperature at high altitude regions remains low, indigenous cold-tolerant bacterium possessing plant growth-promoting (PGP) activities would play an important role in the enhancement of soil nutrients and their utilization by the host plant. Keeping this in consideration, bacterium LJH19 was further evaluated for different PGP attributes (Table 1) (Fig. S3).

Siderophore production by PGPB is also vital for plant defense. Iron chelation by siderophores suppresses fungal pathogens in the rhizosphere (Gulati et al., 2009). In our study, LJH19 was grown in the iron-free CAS broth with pH 6.8 and exhibited considerable siderophore production of 85.72 ±1.06 percent siderophore unit (psu) at 15°C. Further to determine the chemical nature of siderophore, we examined the absorption maxima (□max) of cell-free supernatant in UV-3092 UV/Visible spectrophotometer. We observed a peak at 292nm in the absorption spectra (Fig. S4). In previous study, it was reported that in acidic medium 2,3-dihydroxybenzoic acid (DHB), a phenolic compound consisting of catechol group which absorbs below 330 nm showing two absorption bands with maxima at 254 nm and 292 nm, respectively (Iglesias et al., 2011). DHB is an intermediate involved in the synthesis of catecholate type siderophore (Peralta et al., 2016). These evidences support the presence of DHB in LJH19 grown supernatant indicating the production of catecholate type siderophore.

Production of phytohormone IAA is essential for plant growth to proliferate lateral roots and root hairs (Rosier et al., 2018). In this study, LJH19 demonstrated the ability to produce 166.11 ± 5.7 µg/ml of IAA by colorimetric assay after 72 Hrs of incubation with 200 µg/ml concentration of L-Trytophan at 15°C (Fig. S3A). This infers the ability of LJH19 to produce the IAA in the presence of L-tryptophan signifying the tryptophan dependent pathway of auxin production.

Phosphorous (P) plays an essential role in plant growth and its solubilisation by the action of microorganisms is regarded as the vital PGP trait (Oteino et al., 2015). Since, the plant is unable to uptake inorganic phosphate present in fixed or precipitated form in the soil, bacteria aids in increasing the availability of soluble P for plant acquisition through solubilisation (Santos-Beneit, 2015). Qualitative estimation of phosphate solubilisation showed a 2.3 solubilisation index. Quantitatively, LJH19 was able to solubilise 44.76 ± 1.5 µg/ml of tri-calcium phosphate at 15°C after the 5th day of incubation in NBRIP broth (Fig. S3C). The activity of bacteria was able to decrease pH from 7 to 4.5 indicating the elevation of phosphate solubilisation levels. This suggested that the presence of LJH19 in the compost can deliver available phosphorous to the plants.

Ammonia production by bacteria is yet another feature of PGPB to increase the availability of nitrogen by mineralising organic nitrogen into ammonia (Karthika et al., 2020). In our in-vitro assays, the isolate LJH19 was able to produce a low level of ammonia production (0.20± 0.01 µmoles/ml) (Fig. S3B) after 10 days of incubation in peptone water. These values are quite low in the case of PGP attributes. But, in the case of composting, ammonia gas released by bacteria is primarily responsible for pungent smell and loss of organic nitrogen from the compost (Zhou et al., 2019). This may suggest that it doesn’t directly benefit the plants but may be able to maintain stable organic nitrogen content in the compost by not converting rich nitrogenous sources into ammonia gas.

### Phylogenetic assessment and genome relatedness

A phylogenetic construction based on the complete 16S rRNA gene sequences of a strain belonging to genus *Glutamicibacter* suggests the closest relative of strain LJH19. It was found closest to strain *G. arilaitensis* JB182 but, the type strain of the genus falls in a separate clade (Fig. 2A). The true phylogeny of the isolate was obtained with the phylogenomic tree obtained from PhyloPhlAn, which uses around 400 most conserved gene sequences present across the isolates. Strain LJH19, type strain Re117 and JB182 was found to be in a single clade (Fig. 2B). In order to get the genome relatedness estimate, we have implemented the orthoANI estimation of the isolates from the genus. ANI matrix suggests the genome similarity of the strain LJH19 to subspecies level relatedness to the strain Re117 as its value was around 97% for the type strain Re117 and another strain JB182 (Fig. 2C).

**Fig. 2.**
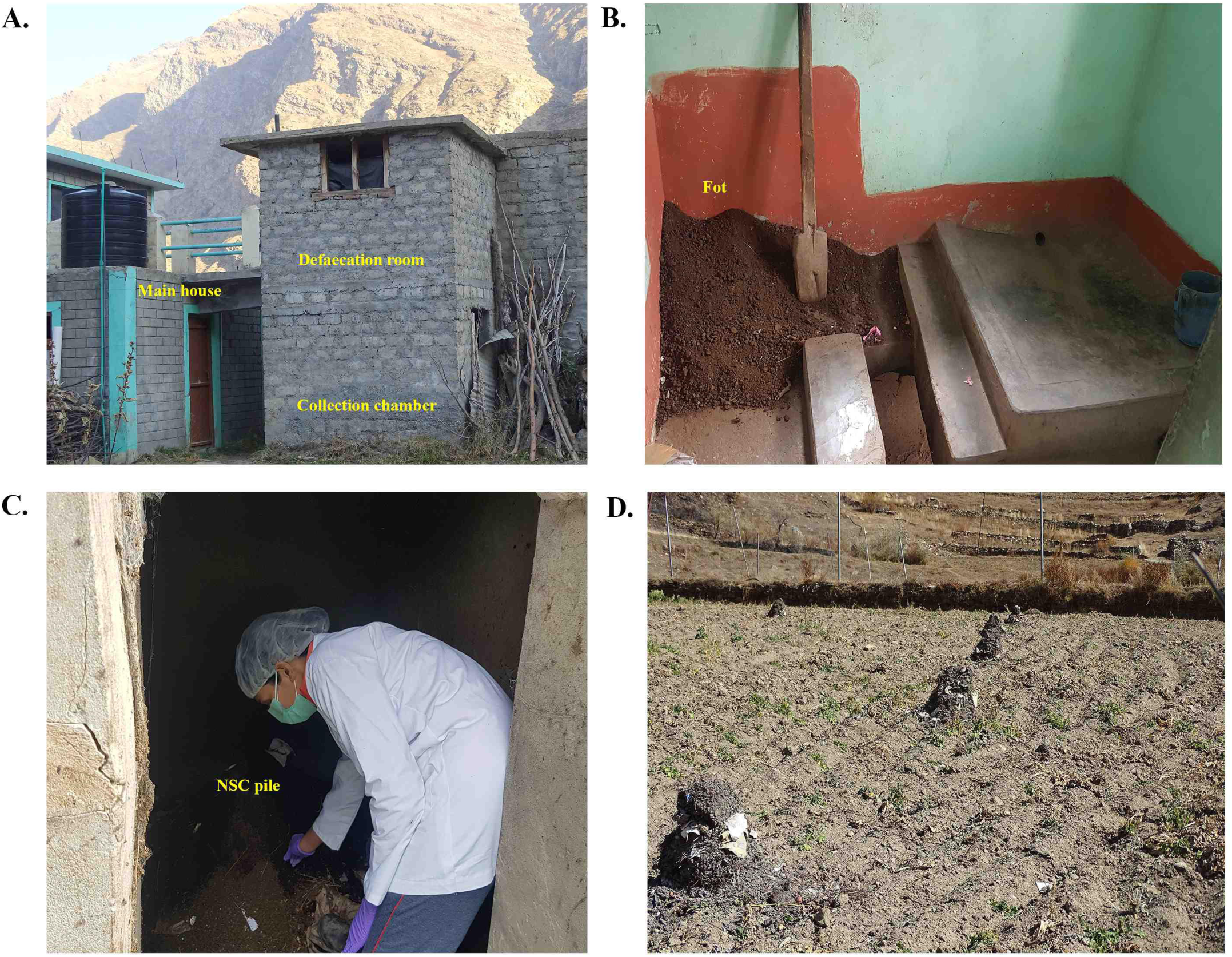
Phylo-taxono-genomics of *Glutamicibacter arilaitensis* LJH19: **A)** 16S rRNA gene phylogeny obtained from the all available *Glutamicibacter* strains. **B)** ML based phylogenomic tree construction obtained from the whole proteome information of the strains of genus *Glutamicibacter*. Violet color circle at each node represents corresponding bootstrap values **C)** OrthoANI similarity matrix created with morpheous, red color represents the maximum values, and yellow color represents the minimum value whereas the green color is the intermediate values. Orange color represents the cutoff value for species demarcation (95% similarity).

### Pan-genome analysis and Chromosomal map

Roary run for the group of the strains forming a clade with the type strain of *Glutamicibacter arilaitensis* and LJH19 resulted in a pan-genome of 9892 genes. A total of 634 genes were found to be core genes, whereas the gene clusters specific to the strain LJH19, Re117 and BJ182 was 1740. A total of 217 genes were specific to the strain LJH19. Chromosomal map showing the unique genomic regions across the strain LJH19 depicts the uniqueness of the strain LJH19 (Fig. 3A). All the strain-specific gene from LJH19 classified by eggNOG falls in several COG categories (Fig. 3B). List of the unique gene, its function and COG classification is reported in Table S2. Based on data retrieved from Prokka annotation and unique genes retrieved from roary run, a representative figure illustrating an overview of different genes involved in catabolic activities, transport, and plant growth promotion in *G. arilaitensis* LJH19 was generated (Fig. 4).

**Fig. 3.**
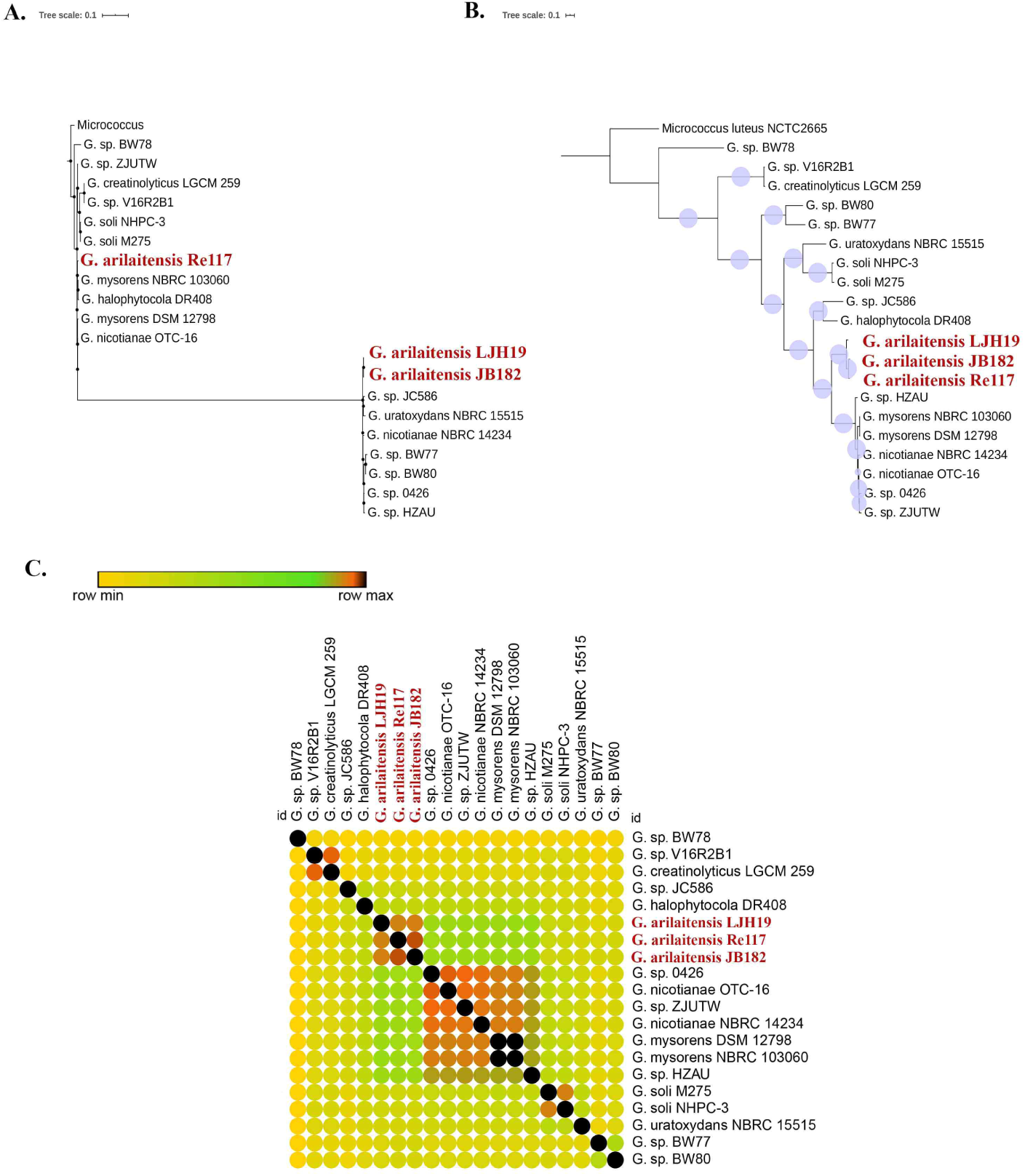
Circular genome representation of LJH19: **A)** BRIG implementation across the three closely related strains including type strain of species Re117, JB182 and LJH19, which resulted in the identification of the unique genomic region across the isolates LJH19. **B)**. Unique genes were fetched from the strain LJH19 using pan genome analysis with the implementation of roary. COG classes and its count identified by eggnog, shows the prevalence of C: Energy production and conversion, E: Amino Acid metabolism and transport, G: Carbohydrate metabolism and transport, I: Lipid metabolism, K: Transcription, L: Replication and repair, M: Cell wall/membrane/envelop biogenesis, and P: Inorganic ion transport and metabolism.

**Fig. 4.**
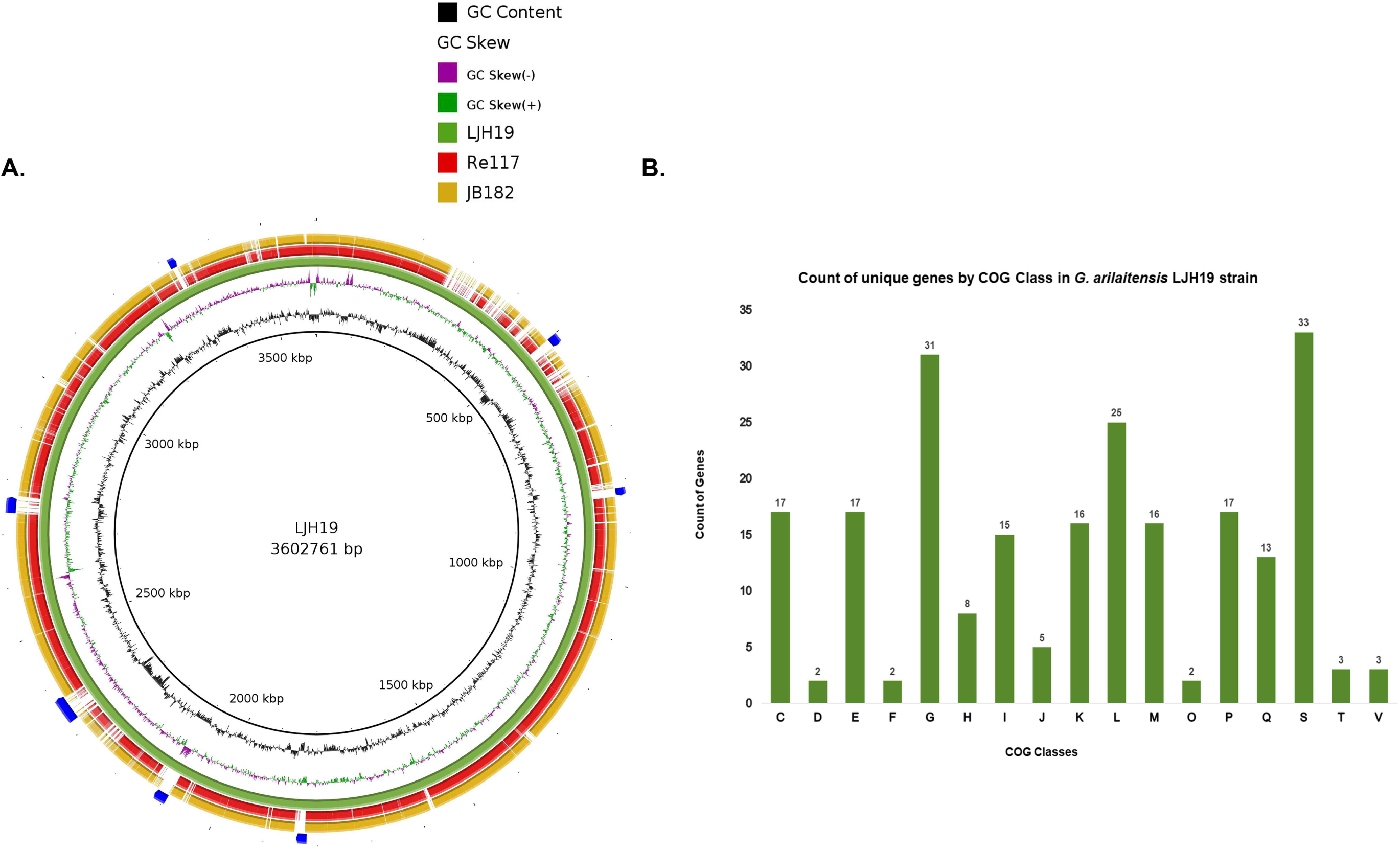
An overview of catabolic activities, transport and plant growth promotion in *G. arilaitensis* LJH19. Following are the selected key genes involved in the pathway : 1, amylase; 2, Oligo-1,6-glucosidase; 3, Beta-glucosidase; 4, Triacylglyceride lipase; 5, monoacylglycerol lipase; 6, Anthranilate synthase component I (TrpE); 7, Anthranilate phosphoribosyl transferase (TrpD); 7,Phosphoribosyl anthranilate isomerase (TrpF); 9, Indole-3-glycerol phosphate synthase (TrpC); 11, Isochorismate synthase (menF); 12, Isochorismatase; 13, Amidase; 14, Argininosuccinate lyase (argH); 15, Arginine decarboxylase (speC); 16, Agmatinase (speB); 17, Polyamine aminopropyltransferase (speE); 18, Ornithine decarboxylase (speC)*. Red arrows indicate enzymes missing in the metabolic pathway. Multistep pathways are denoted with dotted lines. * Gene was annotated as hypothetical protein in Prokka.

### Extended genomic insights on hydrolytic and plant growth promoting attributes of LJH19

The *actinobacteria* are very well known for their ability to produce a variety of secondary metabolites (Dinesh et al., 2017). Hence, we searched for secondary metabolites gene clusters using antiSMASH v5.0 (Blin et al., 2019). This resulted in the identification of three biosynthetic gene cluster namely type III polyketide synthase (PKS), terpene, and siderophore (Fig. 6). The most significant hit predicted by the antiSMASH for type III PKS, terpenes and siderophore is dechlorocuracomycin, carotenoid and desferrioxamin B/desferrioxamine E respectively. Type III PKS are involved in the synthesis of numerous metabolites and have a variety of biological and physiological roles such as defense systems in bacteria (Shimizu et al., 2017). The presence carotenoid gene cluster supports the indicative yellow color of the LJH19 colonies. Besides pigmentation, the major function of carotenoids in bacteria is to protect the cell from UV radiations, oxidative damage and modify membrane fluidity (Liao et al., 2019).

The occurrence of the siderophore gene cluster is demonstrated by the experimental evidence of the *in-vitro* CAS-shuttle assay (Goswami et al., 2014). Through genome mining, we also identified some genes involved in the synthesis of polyamines (PAs), Putrescine (Put) and spermidine (Spd) (TableS2). In bacteria, these active molecules are involved in the biosynthesis of siderophores, improve the survival rate in freezing conditions, and stabilize spheroplasts and protoplasts from osmotic shock (Wortham et al., 2007).

As we discussed earlier, experimental evidence results suggested that LJH19 is involved in the catecholate type siderophore production. The genomic insights further strengthened these findings by predicting the genes involved in the biosynthesis of enterobactin and petrobactin. However, antiSMASH predicted gene clusters for hydroxamate type desferrioxamine B/E siderophore. These results deduce that LJH19 may be able to produce a wide variety of siderophores. Most of the enzymes involved in enterobactin biosynthesis were present except the genes involved in the conversion of 2,3-Dihydro-2,3-dihydroxybenzoate to enterobactin (Fig. 4) (Table S1). We also noticed the genes encoding the transporters required for the import and export of synthesized enterobactin. In respect to the biosynthesis of petrobactin, spermidine molecules are used for synthesis using citrate backbone (Budzikiewicz, 2005). However, no genes were identified in LJH19 for the synthesis of petrobactin. Nevertheless, we identified the genes encoding the transporters required for the import and export of both synthesized enterobactin and petrobactin (Fig. 5) (Table S1). In addition to this, we also found transporters for hydroxamate type siderophores (Table S1). These results correlate with the desferrioxamin B/desferrioxamine E gene cluster identified in antiSMASH run.

**Fig. 5.**
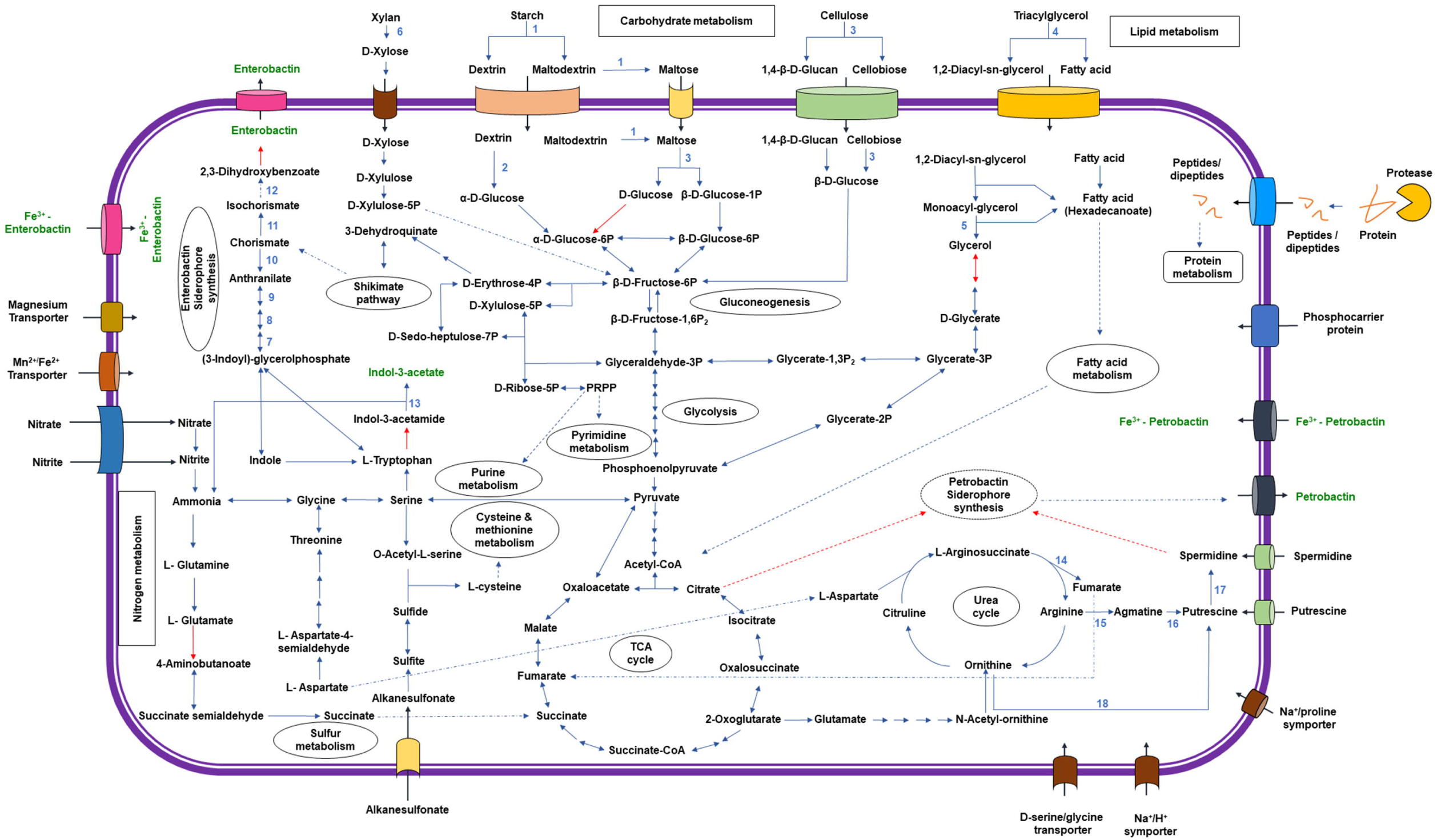
Schematic depiction of unique genes involved in siderophore export, uptake and Ferric-enterobactin processing in LJH19 strain. The genes involved in petrobactin transport are apeX, Apo-petrobactin exporter; fatC/D, Petrobactin import system permease protein; fatE, Petrobactin import ATP-binding protein; yclQ, Petrobactin-binding protein. The genes involved in enterobactin transport are entS, Enterobactin exporter; fepD/G, Ferric enterobactin transport system permease protein; FepC, Ferric enterobactin transport ATP-binding protein; yfiY, putative siderophore-binding lipoprotein. Once translocated inside the bacterial cell, Fe-PB and Fe-EB are further processed by putative esterases to release iron and free PB and EB. Abbreviations are as follows: PUT, putrescine; SPD, spermidine; PB, petrobactin; EB, enterobactin; Fe-PB, ferric bound PB; Fe-EB ferric bound EB. (Hagan et al., 2017, 2016; Krewulak and Vogel, 2016).

**Fig. 6.**
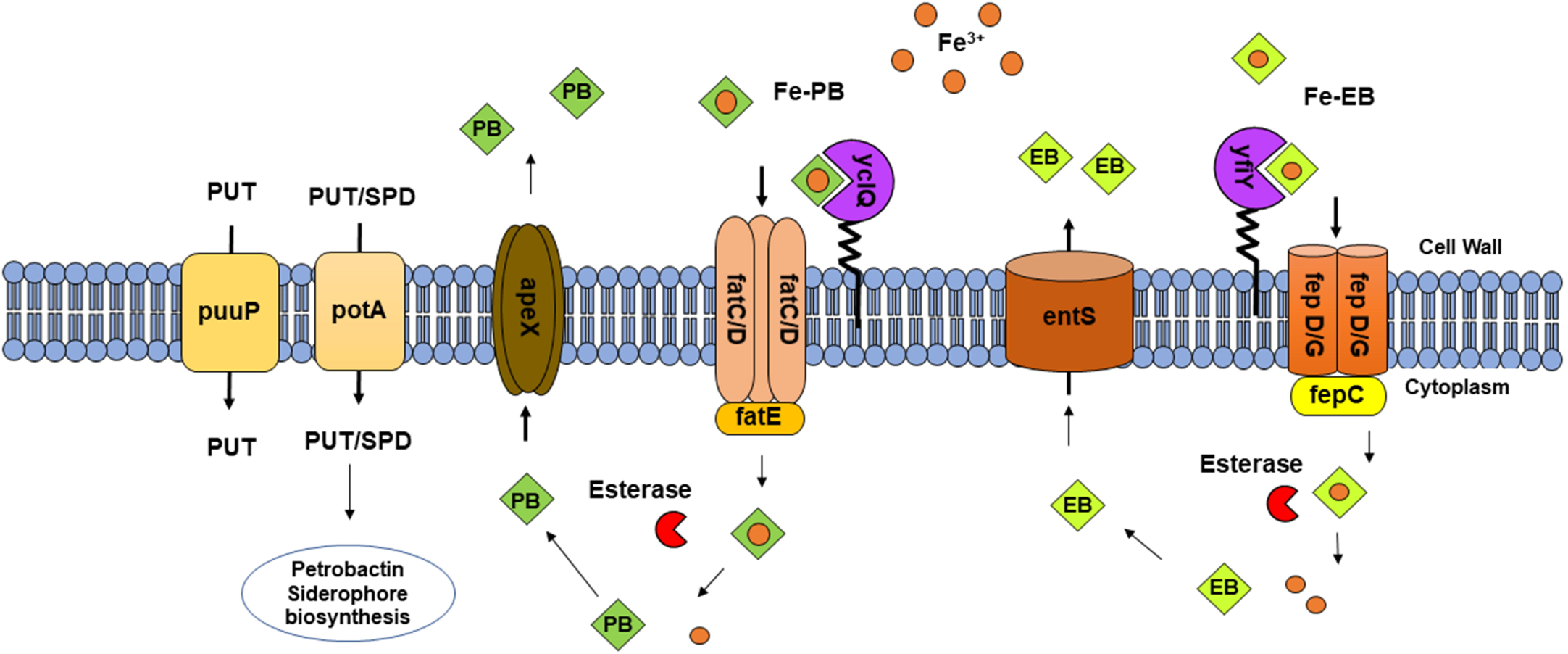
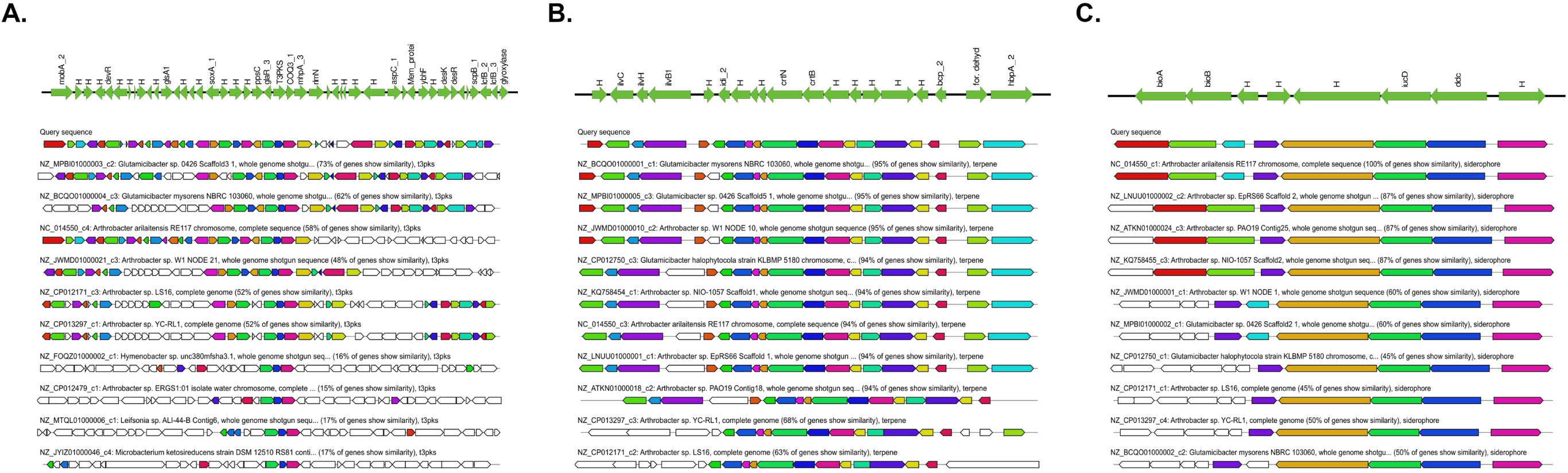
Biosynthetic gene cluster identified by the antiSMASH search namely type III polyketide synthase (PKS), terpene and siderophore. H represents the hypothetical gene cluster annotated by prokka. The direction of arrow represents the forward (5’ → 3’) and reverse orientation of the gene cluster. Among the type III PKS, terpene and siderophore, the predicted gene cluster shows significant hit with the other strains of the genus *Glutamicibacter*. A, B and C represents type III PKS, terpene and siderophore respectively.

A series of genes related to other PGP traits were also identified in the LJH19 genome, the candidate genes that are likely involved in tryptophan-dependent IAA biosynthesis via indole-3-acetamide (IAM) pathway, such as amidase that converts IAM to IAA was present in the genome. However, gene encoding L-tryptophan monooxygenase was missing which converts L-tryptophan to IAM (Li et al., 2018). We also found few genes encoding phosphatases, inositol-phosphatases, and gluconate permease in the genome of LJH19 (Table S1) involved in P metabolism. LJH19 strain has also been noted to carry genes involved in nitrate/nitrite transport pathways including the genes associated with denitrification and nitrate reduction like nitrite reductase and nitrate reductase. Nitrite reductase encoded by the NirD gene converts nitrite to ammonium and further converted to glutamate by glutamate synthetase for amino acid metabolism. These results suggest that LJH19 can deliver plants with available nitrogen sources via enzymatic conversion.

LJH19 strain has several genes encoding proteins involved in the metabolism of a wide variety of complex polysaccharides (Table S1). The depolymerisation of polysaccharides into its oligosaccharides and monomer sugars by bacteria requires a combination of enzymes (Tutino et al., 2002). Many genes in the LJH19 encoded beta-glucosidase, alpha-amylase, beta-xylosidase, pullulanase, oligo-1,6-glucosidase and glycosidases which are involved in the degradation of polysaccharides like cellulose, starch, and xylan (Fig. 4).

The endo-acting enzymes such as alpha-amylase initially hydrolyse the internal linkages of the starch molecule randomly liberating linear and branched oligosaccharides. The pullulanase enzyme specifically cleaves alpha-(1,6)-linkage in pullulan and branched oligosaccharides releasing maltodextrin. Cellulases cleaves the β-(1,4)-glycosidic linkages within the cellulose polymer releasing cellobiose and glucose. Xylan β-(1, 4)-xylosidase cleaves xylan polymers generating smaller oligosaccharides and xylose monomers (Chandra, 2016). Finally, monomeric sugars like glucose and xylose delivered to the cell cytoplasm through specific transporters enter into the glycolysis pathway and ultimately to the TCA cycle generating energy for cellular growth. LJH19 strain have all the crucial genes involved in the core metabolic pathways like glycolysis, citric acid cycle (TCA), gluconeogenesis and pentose phosphate pathway (Fig. 4). LJH19 is also well-equipped with genes encoding proteins that are components of transporter complexes engaged in the recognition and transport of monosaccharide and oligosaccharide such as maltose/maltodextrin, maltooligosaccharide, and cellobiose and transporters for hydrolysed proteins as well (Table S1). Furthermore, we also observed genes such as triacylglycerol lipase which is associated with fatty acid degradation.

LJH19 also harboured several genes related to the adaptational approaches (Table S3). Several cold associated genes encoding for proteins responsible for cold-active chaperons, general stress, osmotic stress, oxidative stress, membrane/cell wall alteration, carbon storage/ starvation, DNA repair, Toxin/Antitoxin modules were identified across the genome. These results infer survival strategies in cold environments.

### *In vitro* and *in silico* pathogenicity analysis of LJH19

LJH19 strain had shown remarkable PGP potential and appreciable hydrolytic activity at low temperature but, to ensure bacterial safety for humans, we determined the pathogenic properties of LJH19 isolate (Table 1). The present study focused on determining the virulence in humans associated with components involved in their colonization into host cells, essential for the commencement of infection. Pathogenic bacteria rely on a variety of virulence factors to induce pathogenesis including adhesion proteins, toxins like haemolysins and proteases (Martínez-García et al., 2018). We performed the initial screening of virulence on blood agar to check the haemolytic activity. In this plate assay, we noticed no haemolytic activity for LJH19 in contrast to haemolytic strains MTCC 96, MTCC121, MTCC 43, MTCC 2470 (Fig. S5A). LJH19 was tested positive for protease activity with an enzymatic index of 12.5 (Fig. S5B), but, quantitatively LJH19 showed very low protease activity (Table 1).

Furthermore, the adherence of bacteria to the host tissue cells is the initial step to induce the pathogenesis (Wilson *et al*., 2019). Therefore, biofilm formation is a notable virulence factor of pathogenic potential. In our in-vitro assay, LJH19 was not able to form biofilm on polystyrene at 37°Cbut, was a weak biofilm producer at 15°C (Fig. S5C). In addition to pathogenesis, biofilm formation also infers antibiotic resistance to the bacterial cells (Patterson et al., 2010). Therefore, we also checked for the resistance phenotype of LJH19, which showed susceptibility to all the twelve antibiotics tested (Fig. S5D), (Table 1).

Virulence is a characteristic of pathogenicity which confers the ability to initiate and sustain infection for the organism. The occurrence of such determinants at the genetic level makes the organism potentially pathogenic with the ability to circulate such genes in the bacterial population (Igbinosa et al., 2017). Since, LJH19 showed some resistance to four antibiotics (Table 1) in our *in-vitro* assays, to affirm the antibiotic resistance obtained from *in-vitro assays* we performed an *insilico* investigation of LJH19 genome and its phylogenetic relative. But, the RGI module of CARD 2020 with strict mode resulted in the detection of no antibiotic resistance gene cluster in LJH19 and its relatives. To further confirm these results, we also assessed the LJH19 genome for its pathogenic potential by PathogenFinder. This web-based tool identifies the genome and provides a probability measure for the test strain to be pathogenic for humans. The predicted results identified LJH19 as a non-human pathogen with an average probability of 0.228 (Supplementary file S4). No putative virulence or pathogenic genes were identified. These results suggested no traces of pathogenicity in LJH19.

## Conclusion

Night soil compost is a rich nutrient source and when supplemented to the soil increases its fertility. In this study, *G. arilaitensis* LJH19 isolated from NSC demonstrated the ability of hydrolysing complex polysaccharides richly found in agricultural residues like starch, cellulose and xylan. The bacterium also exhibited several PGP traits such as IAA production, siderophore production and phosphate solubilization at low ambient temperature. A comprehensive genomic analysis further predicted and excavated some key genes related to the cold adaptation, polysaccharide metabolism, and plant growth promotion. LJH19 also displayed its capabilities as safe bioinoculant by demonstrating negative haemolysis and biofilm formation. Genomic search reinforced the bacterium’s safety with absence of any virulence and antibiotic resistance genes. These results indicated that *G. arilaitensis* LJH19 may serve as a safe bioinoculant and may work collectively in a consortium for efficiently degrading night soil as well as enrich the soil with its PGP attributes. To the best of our knowledge, the current study is the pioneering scientific intervention addressing the issue of NSC in high Himalaya.

## Supporting information

Supplementary output file 1

## Author statement

**Shruti Sinai Borker**: Methodology, Validation, Data curation, Writing-Original draft **Aman Thakur**: Methodology, Writing-Original draft preparation **Sanjeet Kumar**: Software, Writing-Original draft **Sareeka Kumari**: Methodology **Rakshak Kumar**: Conceptualization, Writing – Review & Editing, Supervision, Funding acquisition **Sanjay Kumar**: Project administration

## Acknowledgment

SSB is thankful to UGC, Govt. of India for ‘Research Fellowship’ Grant [UGC-Ref.No.:461/ (CSIR-UGC NET DEC. 2016)]. This work was financially supported by the CSIR in-house FTT project (MLP 0137) and the NMHS project of MoEF&CC (NMHS sanction no. GBPNI/NMHS-2018-19/SG/178). Authors also duly acknowledge the technical support provided by Anil Chaudhary for 16S rRNA gene sequencing, Mohit Kumar Swarnkar for whole genome sequencing and Dr. Vishal Acharya for genome assembly. Authors also acknowledge Aman Kumar for his technical assistance in figure generation. This manuscript represents CSIR-IHBT communication no. **4602**

