## Supplementary figures and images for "Investigation of a night soil compost psychrotrophic bacterium *Glutamicibacter arilaitensis* LJH19 for its safety, efficient hydrolytic and plant growth-promoting potential"

### Supplementary figure S1

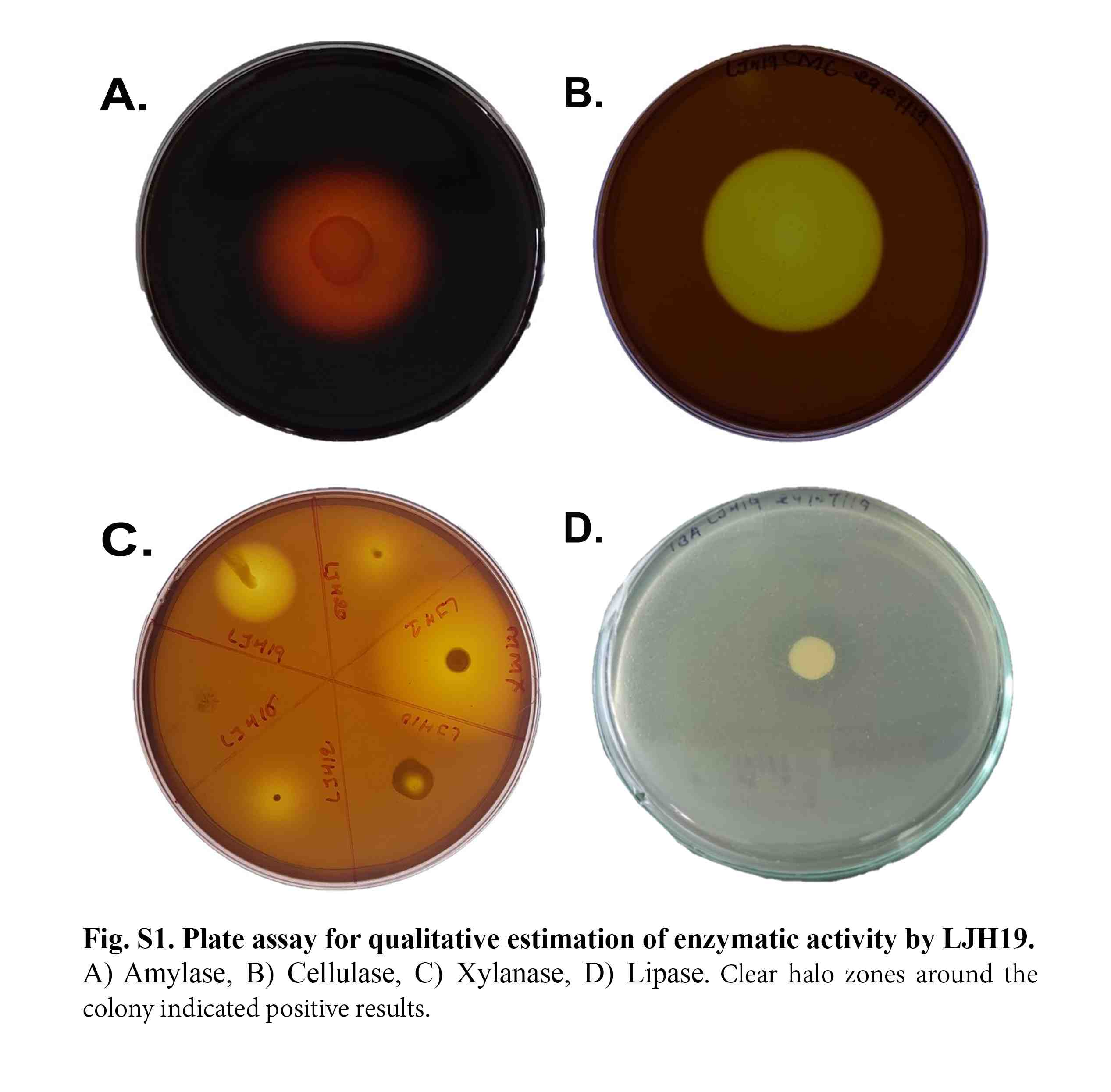

### Supplementary figure S2

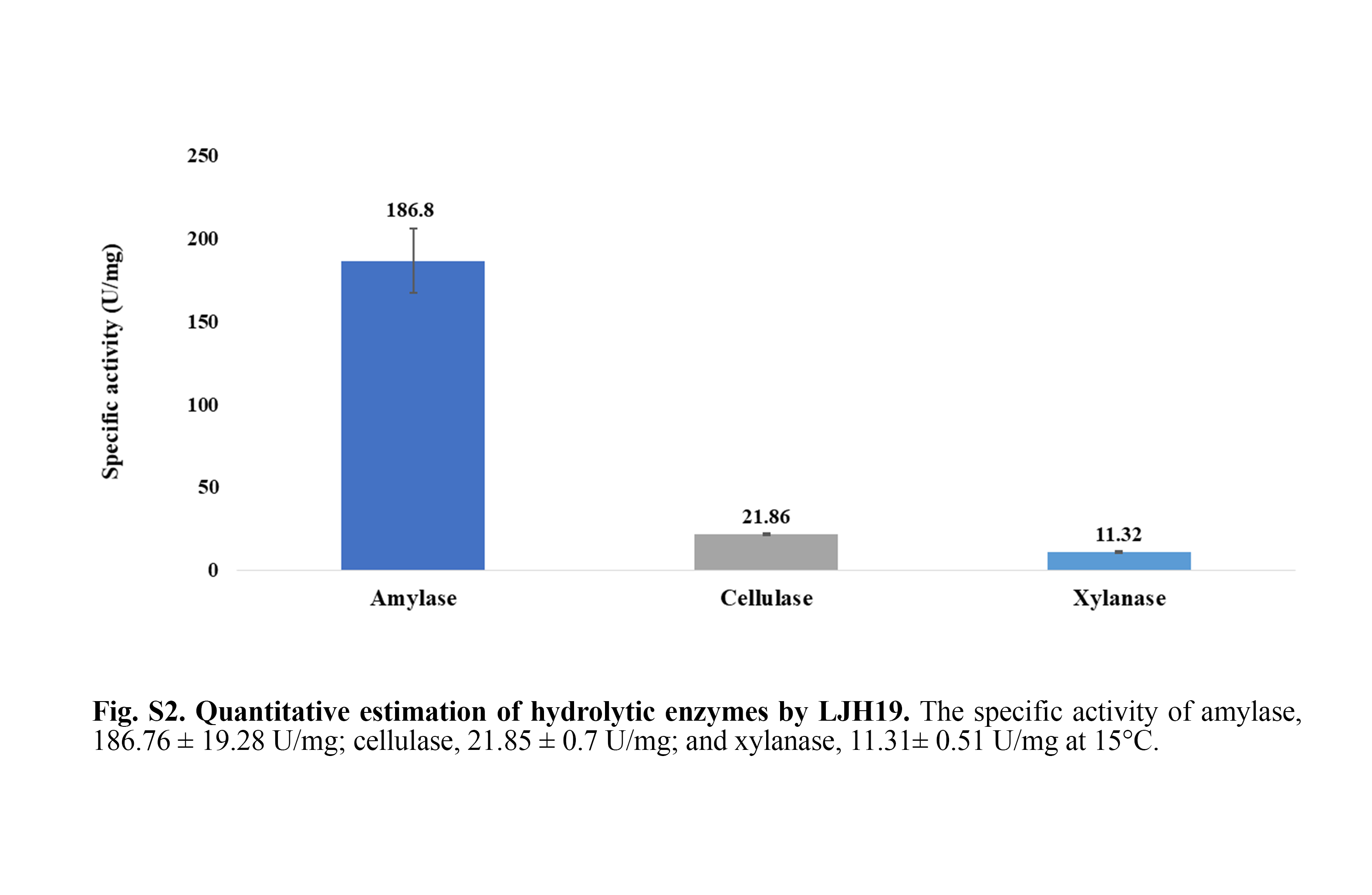

### Supplementary figure S3

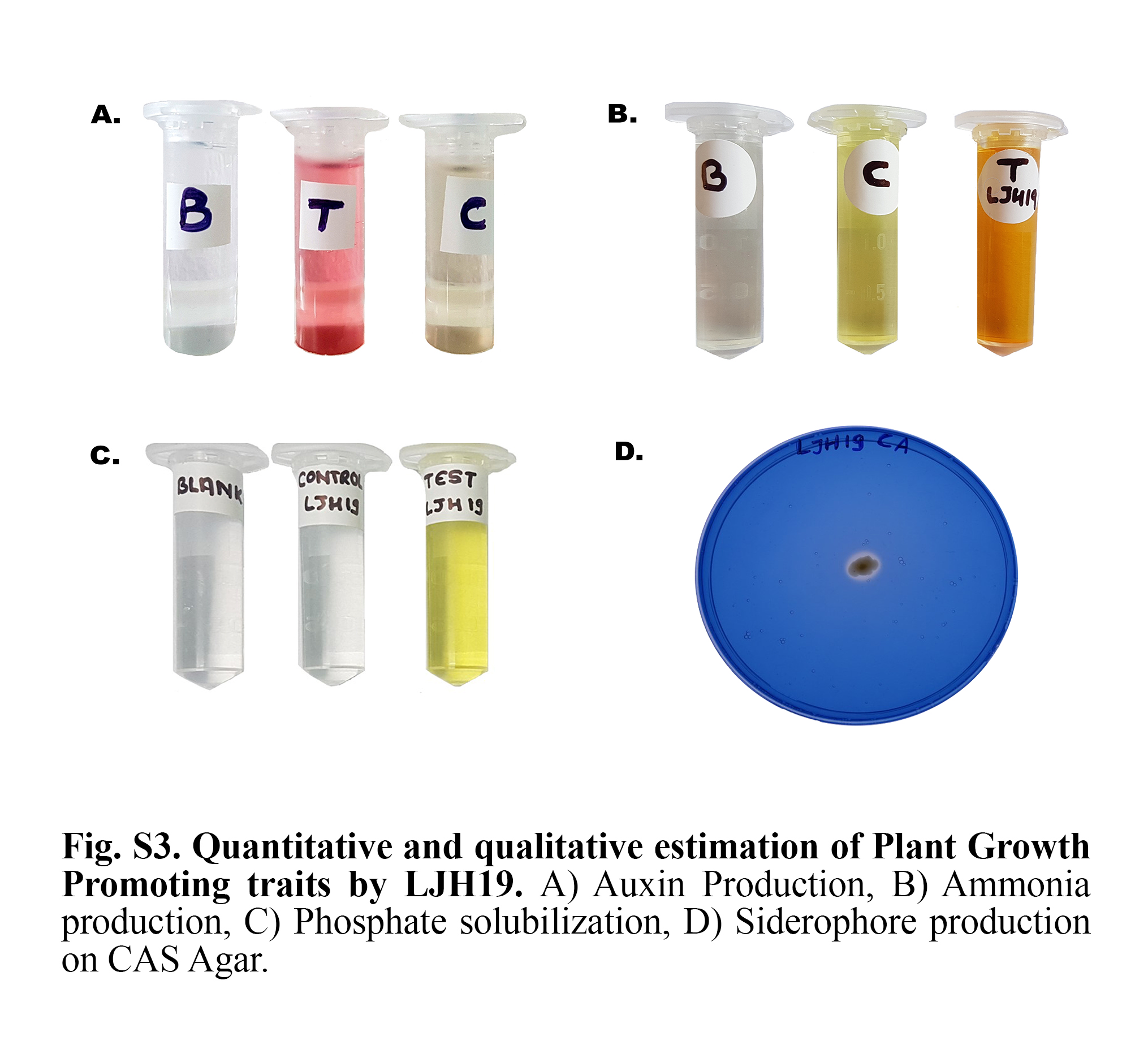

### Supplementary figure S4

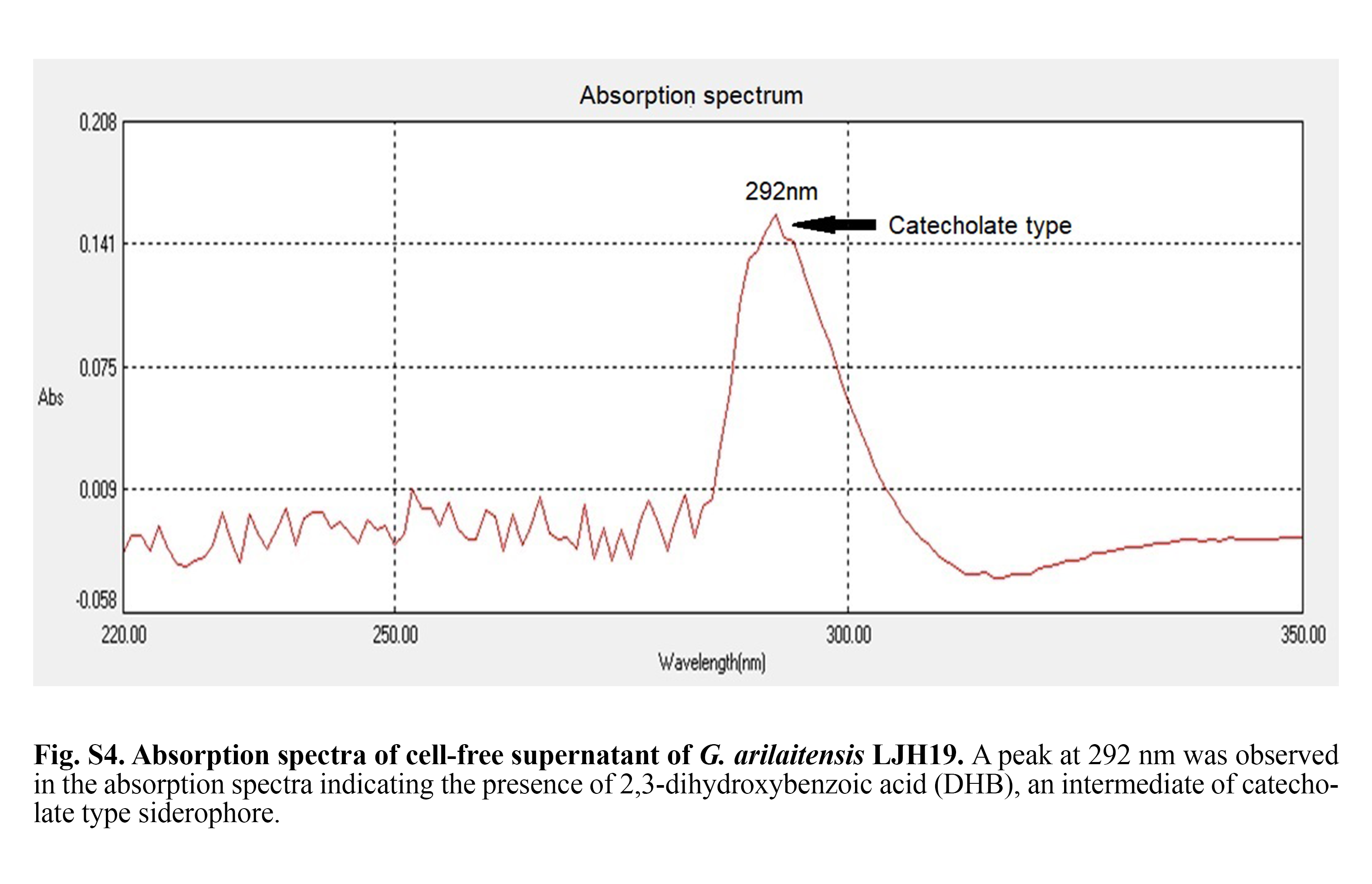

### Supplementary figure S5

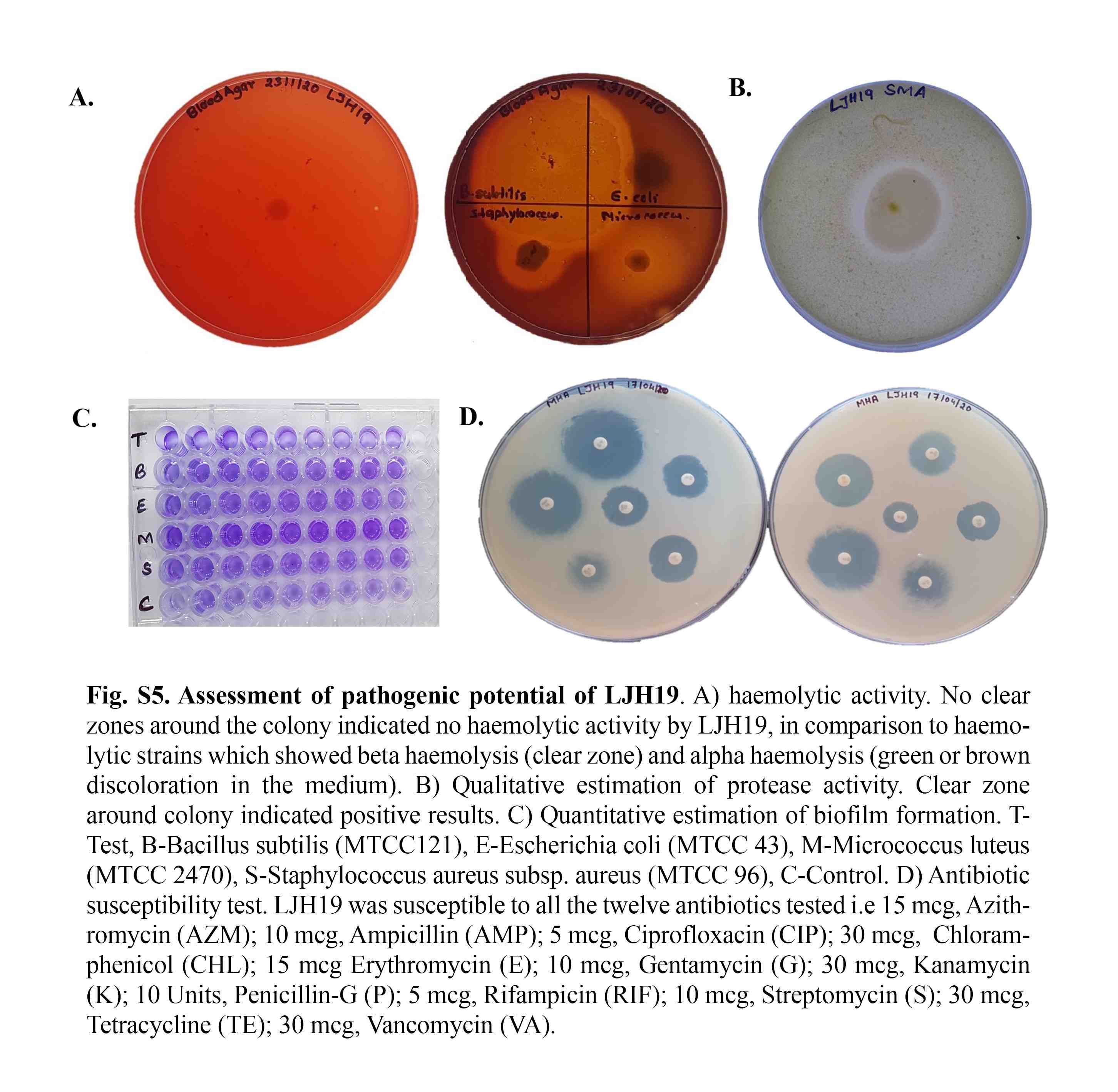
