## Supplementary Table 1 for "Investigation of a night soil compost psychrotrophic bacterium *Glutamicibacter arilaitensis* LJH19 for its safety, efficient hydrolytic and plant growth-promoting potential"

**Supplementary Table 1: List of genes involved in catabolic activity, transport, and PGP activity**

| **Locus_tag** | **gene** | **EC_number** | **COG** | **Product** |
| --- | --- | --- | --- | --- |
| **Catabolic activity** |  |  |  |  |
| MJJNNIGE_00069 | pepO | 3.4.24.- | COG3590 | Neutral endopeptidase |
| MJJNNIGE_00216 |  | 3.1.2.- | COG2050 | Putative esterase |
| MJJNNIGE_00245 | lacZ | 3.2.1.23 |  | Beta-galactosidase |
| MJJNNIGE_00546 |  | 3.2.1.- |  | Beta-xylosidase |
| MJJNNIGE_01053 | pulA | 3.2.1.41 |  | Pullulanase |
| MJJNNIGE_01084 |  | 3.4.21.- | COG0265 | Serine protease |
| MJJNNIGE_01136 | aml | 3.2.1.1 |  | Alpha-amylase |
| MJJNNIGE_01285 | sacC | 3.2.1.80 | COG1621 | Levanase |
| MJJNNIGE_01544,  MJJNNIGE_01174 | htpX | 3.4.24.- |  | Protease HtpX |
| MJJNNIGE_02347 | malL | 3.2.1.10 | COG0366 | Oligo-1,6-glucosidase |
| MJJNNIGE_02452 |  | 3.1.1.23 |  | Thermostable monoacylglycerol lipase |
| MJJNNIGE_02932 | rip1 | 3.4.24.- | COG0750 | Zinc metalloprotease Rip1 |
| MJJNNIGE_03107 | caeB | 3.1.1.- |  | Carboxylesterase B |
| MJJNNIGE_03134 | bglA | 3.2.1.21 | COG2723 | Beta-glucosidase A |
| MJJNNIGE_03364 | xylA | 5.3.1.5 |  | Xylose isomerase |
| MJJNNIGE_03385 |  | 3.2.1.- |  | Beta-xylosidase |
| **Plant growth promoting activity** |  |  |  |  |
| MJJNNIGE_00110 | amiB2 | 3.5.1.4 | COG0154 | Putative amidase AmiB2 |
| MJJNNIGE_00146 | trpG | 4.1.3.27 | COG0512 | Anthranilate synthase component 2 |
| MJJNNIGE_00383 | hpaH | 1.14.14.8 | COG2368 | Anthranilate 3-monooxygenase oxygenase component |
| MJJNNIGE_00674,  MJJNNIGE_00678 | nasD | 1.7.1.4 | COG1251 | Nitrite reductase [NAD(P)H] |
| MJJNNIGE_00675 |  |  |  | hypothetical protein |
| MJJNNIGE_00679 | nasC | 1.7.-.- | COG0243 | Assimilatory nitrate reductase catalytic subunit |
| MJJNNIGE_00796,  MJJNNIGE_00881 | andAa | 1.18.1.3 |  | Anthranilate 1,2-dioxygenase system ferredoxin--NAD(+) reductase component |
| MJJNNIGE_00805 | puo | 1.4.3.10 |  | Putrescine oxidase |
| MJJNNIGE_01047,  MJJNNIGE_01165 | menF | 5.4.4.2 |  | Isochorismate synthase MenF |
| MJJNNIGE_01620 | trpE | 4.1.3.27 | COG0147 | Anthranilate synthase component 1 |
| MJJNNIGE_01728 | sir | 1.8.7.1 | COG0155 | Sulfite reductase [ferredoxin] |
| MJJNNIGE_01926 | trpD | 2.4.2.18 | COG0547 | Anthranilate phosphoribosyltransferase |
| MJJNNIGE_02216, MJJNNIGE_03410 | speE | 2.5.1.16 |  | Polyamine aminopropyltransferase (Spermidine synthase) |
| MJJNNIGE_03196 | yecD | 3.-.-.- | COG1335 | Isochorismatase family protein YecD |
| MJJNNIGE_03076 | argD | 2.6.1.11 | COG4992 | Acetylornithine aminotransferase |
| MJJNNIGE_03077 | argF | 2.1.3.3 | COG0078 | Ornithine carbamoyltransferase |
| MJJNNIGE_03081 | argH | 4.3.2.1 | COG0165 | Argininosuccinate lyase |
| MJJNNIGE_00820 | hutG | 3.5.3.8 | COG0010 | Formimidoylglutamase* (Arginase) |
| MJJNNIGE_00897 | speC |  |  | hypothetical protein*(ornithine decarboxylase) |
| MJJNNIGE_02562 | speB |  |  | Guanidinobutyrase (agmatinase) |
| **Transporters** |  |  |  |  |
| MJJNNIGE_00015,  MJJNNIGE_00589,  MJJNNIGE_00968, | yhdG |  | COG0531 | putative amino acid permease YhdG |
| MJJNNIGE_00049 | pstB1 | 7.3.2.1 | COG1117 | Phosphate import ATP-binding protein PstB 1 |
| MJJNNIGE_00050 |  |  |  | hypothetical protein |
| MJJNNIGE_00051 | pstC2 |  | COG0573 | Phosphate transport system permease protein PstC 2 |
| MJJNNIGE_00122 | xylE |  |  | D-xylose-proton symporter |
| MJJNNIGE_00111 | dtpT |  | COG3104 | Di-/tripeptide transporter |
| MJJNNIGE_00242 | xylG | 7.5.2.10 | COG1129 | Xylose import ATP-binding protein XylG |
| MJJNNIGE_00243 | xylH |  | COG4214 | Xylose transport system permease protein XylH |
| MJJNNIGE_00373, MJJNNIGE_00374,  MJJNNIGE_03335 | ssuC |  | COG0600 | Putative aliphatic sulfonates transport permease protein SsuC |
| MJJNNIGE_00413 | fatE | 7.2.2.- | COG4604 | Petrobactin import ATP-binding protein FatE |
| MJJNNIGE_00414 | fatC |  | COG4605 | Petrobactin import system permease protein FatC |
| MJJNNIGE_00415 | fatD |  | COG4606 | Petrobactin import system permease protein FatD |
| MJJNNIGE_00416 | yclQ |  | COG4607 | Petrobactin-binding protein YclQ |
| MJJNNIGE_01051, MJJNNIGE_01739 | apeX |  | COG2409 | Apo-petrobactin exporter |
| MJJNNIGE_00510 | puuP |  | COG0531 | Putrescine importer PuuP |
| MJJNNIGE_00547,  MJJNNIGE_00784,  MJJNNIGE_03386 | ngcG |  | COG0395 | Diacetylchitobiose uptake system permease protein NgcG |
| MJJNNIGE_00548 | ngcF |  | COG1175 | Diacetylchitobiose uptake system permease protein NgcF |
| MJJNNIGE_00551 |  |  | COG1132 | putative ABC transporter ATP-binding protein |
| MJJNNIGE_00552 |  | 7.6.2.- | COG1132 | Fatty acid ABC transporter ATP-binding/permease protein |
| MJJNNIGE_00597 | cycA |  | COG1113 | D-serine/D-alanine/glycine transporter |
| MJJNNIGE_00680 | narK |  | COG2223 | Nitrate/nitrite transporter NarK |
| MJJNNIGE_00779 | gntT |  | COG2610 | High-affinity gluconate transporter |
| MJJNNIGE_00959 | yfiZ |  | COG0609 | putative siderophore transport system permease protein YfiZ |
| MJJNNIGE_00960 | yfhA |  | COG0609 | putative siderophore transport system permease protein YfhA |
| MJJNNIGE_01002 | potD |  | COG0687 | Spermidine/putrescine-binding periplasmic protein |
| MJJNNIGE_01004 | potA | 7.6.2.11 | COG3842 | Spermidine/putrescine import ATP-binding protein PotA |
| MJJNNIGE_01022, MJJNNIGE_02404,  MJJNNIGE_02770,  MJJNNIGE_03351 | entS |  |  | Enterobactin exporter EntS |
| MJJNNIGE_01290,  MJJNNIGE_01459,  MJJNNIGE_02914, | dppB |  | COG0601 | Dipeptide transport system permease protein DppB |
| MJJNNIGE_01291 | dppC |  | COG1173 | Dipeptide transport system permease protein DppC |
| MJJNNIGE_01292 | dppD |  | COG0444 | Dipeptide transport ATP-binding protein DppD |
| MJJNNIGE_01293 | oppF |  | COG4608 | Oligopeptide transport ATP-binding protein OppF |
| MJJNNIGE_01734 | nrtB |  | COG0600 | Nitrate import permease protein NrtB |
| MJJNNIGE_02149 | yfhA_2 |  | COG0609 | putative siderophore transport system permease protein YfhA |
| MJJNNIGE_02150 | yfiZ_2 |  | COG0609 | putative siderophore transport system permease protein YfiZ |
| MJJNNIGE_02151 | yfiY |  | COG0614 | putative siderophore-binding lipoprotein YfiY |
| MJJNNIGE_02344 | malG |  | COG3833 | Maltose/maltodextrin transport system permease protein MalG |
| MJJNNIGE_02345 | malF |  | COG1175 | Maltose/maltodextrin transport system permease protein MalF |
| MJJNNIGE_02346 | cycB |  | COG2182 | Cyclodextrin-binding protein |
| MJJNNIGE_02833 | melC |  | COG0395 | Melibiose/raffinose/stachyose import permease protein MelC |
| MJJNNIGE_02856 | fepC |  | COG1120 | Ferric enterobactin transport ATP-binding protein FepC |
| MJJNNIGE_02857 | fepG |  | COG4779 | Ferric enterobactin transport system permease protein FepG |
| MJJNNIGE_02858 | fepD |  | COG0609 | Ferric enterobactin transport system permease protein FepD |
| MJJNNIGE_02913 | oppA |  | COG4166 | Oligopeptide-binding protein OppA |
| MJJNNIGE_02915 | oppC |  | COG1173 | Oligopeptide transport system permease protein OppC |
| MJJNNIGE_02920 |  |  | COG1173 | Putative peptide transport permease protein |
| MJJNNIGE_03304 | fepB |  | COG4592 | Ferrienterobactin-binding periplasmic protein |
| MJJNNIGE_03333 | nrtD | 7.3.2.4 | COG1116 | Nitrate import ATP-binding protein NrtD |
| MJJNNIGE_03387 | melD |  | COG1175 | Melibiose/raffinose/stachyose import permease protein MelD |
