## Supplementary Table 3 for "Investigation of a night soil compost psychrotrophic bacterium *Glutamicibacter arilaitensis* LJH19 for its safety, efficient hydrolytic and plant growth-promoting potential"

**Supplementary Table 3.** Genes encoding known cold & stress response and DNA repair proteins as predicted in the genome of *G. arilaitensis* LJH19**.**

| **Locus_tag** | **Gene** | **COG** | **Product** |
| --- | --- | --- | --- |
| **Cold active chaperones** |  |  |  |
| MJJNNIGE_00922, MJJNNIGE_01807 | cspA | COG1278 | Cold shock-like protein CspA |
| MJJNNIGE_01301,  MJJNNIGE_02533 | groL1 | COG0459 | 60 kDa chaperonin 1 |
| MJJNNIGE_01300 | groS | COG0234 | 10 kDa chaperonin |
| MJJNNIGE_01058, MJJNNIGE_01444, MJJNNIGE_01900 | dnaJ | COG0484 | Chaperone protein DnaJ |
| MJJNNIGE_01056 | dnaK | COG0443 | Chaperone protein DnaK |
| MJJNNIGE_01067 | clpB | COG0542 | Chaperone protein ClpB |
| MJJNNIGE_01896 | hemW | COG0635 | Heme chaperone HemW |
| MJJNNIGE_00929 | hslR | COG1188 | Heat shock protein 15 |
| **Osmotic Stress/ Oxidative stress** |  |  |  |
| MJJNNIGE_03015 | opuCB | COG1174 | Glycine betaine/carnitine/choline transport system permease protein OpuCB |
| MJJNNIGE_03016 | opuCC | COG1732 | Glycine betaine/carnitine/choline-binding protein OpuCC |
| MJJNNIGE_00456, MJJNNIGE_03337 | betP | COG1292 | Glycine betaine transporter BetP |
| MJJNNIGE_00873 | soxB |  | Sarcosine oxidase subunit beta |
| MJJNNIGE_00874 | soxD |  | Sarcosine oxidase subunit delta |
| MJJNNIGE_00626, MJJNNIGE_00875 | soxA |  | Sarcosine oxidase subunit alpha |
| MJJNNIGE_00876 | soxG |  | Sarcosine oxidase subunit gamma |
| MJJNNIGE_01666 | sodA | COG0605 | Superoxide dismutase [Mn] |
| MJJNNIGE_02183 | katE | COG0753 | Catalase HPII |
| MJJNNIGE_02673 | katA | COG0753 | Catalase |
| MJJNNIGE_03339, MJJNNIGE_03340 | katG | COG0376 | Catalase-peroxidase |
| MJJNNIGE_02183 | katE | COG0753 | Catalase HPII |
| MJJNNIGE_00518, MJJNNIGE_02296 | bcp | COG1225 | Putative peroxiredoxin |
| MJJNNIGE_02122 | ahpE | COG1225 | Alkyl hydroperoxide reductase E |
| MJJNNIGE_00020 | ohrB | COG1764 | Organic hydroperoxide resistance protein OhrB |
| MJJNNIGE_00224 | ohrR |  | Organic hydroperoxide resistance transcriptional regulator |
| MJJNNIGE_00026, MJJNNIGE_01178 | trxC | COG0526 | Putative thioredoxin 2 |
| MJJNNIGE_00179 | trxA |  | Thioredoxin 1 |
| MJJNNIGE_00180 | trxB | COG0492 | Thioredoxin reductase |
| MJJNNIGE_01723 |  |  | Thioredoxin-like reductase |
| MJJNNIGE_02757 | nhaA | COG3004 | Na(+)/H(+) antiporter NhaA |
| MJJNNIGE_01038 | mrpD | COG0651 | Na(+)/H(+) antiporter subunit D |
| MJJNNIGE_01041 | mrpG | COG1320 | Na(+)/H(+) antiporter subunit G |
| MJJNNIGE_02580 | czcD | COG1230 | Cadmium, cobalt and zinc/H(+)-K(+) antiporter |
| MJJNNIGE_02518 | otsB | COG0637 | Trehalose-phosphate phosphatase |
| MJJNNIGE_02519 | otsA | COG0380 | Trehalose-6-phosphate synthase |
| **General Stress response** |  |  |  |
| MJJNNIGE_00131, MJJNNIGE_00443, MJJNNIGE_00778, MJJNNIGE_00806, MJJNNIGE_02977, MJJNNIGE_03407 |  | COG0589 | Universal stress protein |
| MJJNNIGE_00144, MJJNNIGE_02449, MJJNNIGE_02788 | pknD |  | Serine/threonine-protein kinase PknD |
| MJJNNIGE_00075 | hipA | COG3550 | Serine/threonine-protein kinase toxin HipA |
| MJJNNIGE_00145 | pknB | COG0515 | Serine/threonine-protein kinase PknB |
| MJJNNIGE_00020 | ohrB | COG1764 | Organic hydroperoxide resistance protein OhrB |
| MJJNNIGE_00224 | ohrR |  | Organic hydroperoxide resistance transcriptional regulator |
| MJJNNIGE_02122 | ahpE | COG1225 | Alkyl hydroperoxide reductase E |
| MJJNNIGE_02752 | rplY | COG1825 | 50S ribosomal protein L25 |
| MJJNNIGE_01814 | yedK | COG2135 | SOS response-associated protein YedK |
| **Carbon storage/starvation** |  |  |  |
| MJJNNIGE_00842 | dps | COG0783 | DNA protection during starvation protein |
| MJJNNIGE_01661 | plsC | COG0204 | 1-acyl-sn-glycerol-3-phosphate acyltransferase |
| **DNA repair** |  |  |  |
| MJJNNIGE_02021, MJJNNIGE_02131 | xerC |  | Tyrosine recombinase XerC |
| MJJNNIGE_02490 | xerD | COG4974 | Tyrosine recombinase XerD |
| MJJNNIGE_03031 | recA | COG0468 | Protein RecA |
| MJJNNIGE_02061 | dnaC | COG0305 | Replicative DNA helicase |
| MJJNNIGE_02218 | recG |  | ATP-dependent DNA helicase RecG |
| MJJNNIGE_02550, MJJNNIGE_01428 | uvrD1 | COG0210 | ATP-dependent DNA helicase UvrD1 |
| MJJNNIGE_01424, MJJNNIGE_01425 | rep |  | ATP-dependent DNA helicase Rep |
| MJJNNIGE_01838 | ruvA | COG0632 | Holliday junction ATP-dependent DNA helicase RuvA |
| MJJNNIGE_01839 | ruvB | COG2255 | Holliday junction ATP-dependent DNA helicase RuvB |
| MJJNNIGE_00520 | ligD | COG1793 | Multifunctional non-homologous end joining DNA repair protein LigD |
| MJJNNIGE_01911 | recO |  | DNA repair protein RecO |
| MJJNNIGE_02493 | recN | COG0497 | DNA repair protein RecN |
| MJJNNIGE_00054 | radA | COG1066 | DNA repair protein RadA |
| **Toxin/Antitoxin modules** |  |  |  |
| MJJNNIGE_00212 | higA2 |  | Putative antitoxin HigA2 |
| MJJNNIGE_00075 | hipA | COG3550 | Serine/threonine-protein kinase toxin HipA |
